# Severe Hyposmia Distinguishes Neuropathologically Confirmed Dementia with Lewy Bodies from Alzheimer’s Disease Dementia

**DOI:** 10.1101/743856

**Authors:** TG Beach, CH Adler, N Zhang, GE Serrano, LI Sue, Erika Driver-Dunckley, Shayamal H. Mehta, E Zamrini, MN Sabbagh, HA Shill, CM Belden, DR Shprecher, RJ Caselli, EM Reiman, KJ Davis, KE Long, LR Nicholson, AJ Intorcia, MJ Glass, JE Walker, M Callan, JC Oliver, R Arce, RC Gerkin

## Abstract

Due to the absence of core clinical features, many subjects with neuropathologically-confirmed dementia with Lewy bodies (DLB) are never diagnosed as such during life. Most of these are diagnosed with Alzheimer’s disease dementia (ADD) or unspecified dementia. Unrecognized DLB therefore is a critical impediment to clinical studies and treatment trials of both ADD and DLB. There are numerous published studies that suggest that olfactory function tests may be able to differentiate some neurodegenerative conditions from each other and from normal subjects, but there are very few studies with neuropathological confirmation of diagnosis. We compared University of Pennsylvania Smell Identification Test (UPSIT) results in 209 subjects: 1) 29 concurrently meeting intermediate or high consensus clinicopathological criteria for both DLB and ADD 2) 96 meeting criteria for ADD without DLB 3) 84 control subjects that were non-demented and without parkinsonism at death. The DLB subjects had significantly lower (one-way ANOVA p < 0.0001, pairwise Bonferroni p < 0.05) first and mean UPSIT scores (13.7 and 13.2) than ADD (23.3 and 22.2) or controls (29.6 and 28.9). For subjects with first and mean UPSIT scores less than 20 and 17, respectively, Firth logistic regression analysis, adjusted for age, gender and mean MMSE score, conferred statistically significant odds ratios of 17.5 and 18.0 for predicting a DLB vs ADD diagnosis, as compared to 3.3 for the presence or absence of visual hallucinations throughout the clinical observation period. To our knowledge, this is the largest study to date comparing olfactory function in subjects with neuropathologically confirmed DLB and ADD. Olfactory function testing may be a convenient and inexpensive strategy for enriching dementia studies or clinical trials with DLB subjects, or conversely, reducing the inclusion of DLB subjects in ADD studies or trials.

## Introduction

Dementia due to AD (ADD) is often associated with comorbid brain disease that may affect clinical presentation, rate of cognitive decline, and response to therapeutic agents [1–22]. Additional concurrent pathology could be unresponsive to therapies directed at the “primary” pathology. It is apparent then, that clinical trials for ADD could suffer from decreased effect size if this were true.

The most common comorbidity in ADD is Lewy body disease. Slightly more than one-half or more of all those meeting clinicopathological ADD diagnostic criteria also have α-synuclein pathology [9,23–25] with morphological features similar to Parkinson’s disease (PD). This is broadly termed “Lewy body disease” (LBD). Similarly, about one-half of those with dementia and PD (PDD) [26–38] and three-quarters or more of those with dementia with Lewy bodies (DLB), have clinically significant AD histopathology [39–42]. In the majority of subjects with ADD and DLB, the typical clinical signs and symptoms of DLB [43,44] are absent and thus this co-existence is recognized only at autopsy [22,45–47]. This clinical inability to separate ADD from DLB hampers clinical trials for both conditions. Several autopsy-validated studies have indicated that cognitive decline is faster in elderly subjects dying with ADD who also have LBD [3,22,48–51], and disease duration has been reported to be shorter in those with coexistent ADD and DLB [40,52]. There is therefore a critical need for better clinical differentiation of these two conditions.

There are numerous published clinical studies that suggest that olfactory function tests may be useful in differentiating amongst cerebrovascular and neurodegenerative disorders [53–64] and, in particular, in distinguishing DLB from ADD [65–70], but the studies with later neuropathological establishment of the specific molecular pathology are the most informative. Possibly the first such study, done by Oxford University [71], investigated the neuropathological correlates of anosmia in subjects with dementia. Anosmia was defined on the basis of being able or unable to detect the scent of lavender oil. Seventeen subjects had neuropathological DLB, defined as the concurrent presence of Lewy bodies in both the substantia nigra and cingulate gyrus. Sixteen of these had concurrent ADD while another 43 subjects had ADD alone, defined as probable or definite CERAD AD [72], without LBD. Anosmia was significantly (p = 0.029) more common in DLB (41%) than in ADD (16%).

A similar study [73] from the University of Southern California defined anosmia as the inability to detect the odor of N-butyl alcohol, finding anosmia in 47% of those with the Lewy body variant (LBV, n = 17)) of AD versus 22.% of those with AD alone (n = 89). This proportional difference was highly significant (p = 0.0004). The diagnosis of LBV was defined as the presence of Lewy bodies in both brainstem and cerebral cortex while ADD was defined as CERAD probable or definite AD. The independent odds ratio for anosmia as a predictor of LBV was 5.4 vs 7.3 for visual hallucinations.

In a study of a mixed group of non-demented and demented subjects with and without parkinsonism from the Rush Memory and Aging Project [74], lower scores on the Brief Smell Identification Test were significant predictors of limbic and neocortical LBD stages (9 and 13 subjects, respectively). The presence of any Lewy bodies accounted for 15.4% of test variance, as compared to 4.1% due to a composite measure of AD histopathology.

Incidental Lewy body disease (ILBD) refers to the presence of LBD in asymptomatic elderly people and is likely to be a prodromal stage of PD or DLB as striatal dopaminergic markers are halfway between asymptomatic elderly people without LBD and clinically-manifest PD [75–78]. One neuropathologically-informed study has reported that olfactory function in subjects with ILBD is also halfway between PD and asymptomatic elderly people without LBD, suggesting that hyposmia may be useful as a prodromal marker [79]. Another prior study found an OR of 11.0 for the prediction of ILBD in those amongst the lowest tertile of olfactory function [80]. Supporting the possible usefulness of hyposmia as a prodromal biomarker are the findings that it is present in some clinically normal GBA and LRRK2 mutation carriers [81,82], is common in idiopathic REM sleep behavior disorder (iRBD) [83–85], is a significant and independent predictor of phenoconversion from iRBD to parkinsonism or dementia [86] and is associated with decreased striatal dopamine transporter imaging [87,88]. A likely causative factor underlying the impairment of olfaction in LBD is the near-universal occurrence of early-stage α-synuclein pathology within the olfactory bulb [89–92].

In this study we sought to determine the diagnostic utility of hyposmia as a diagnostic predictor of neuropathologically-identified DLB, using the largest set to date of neuropathologically-examined subjects.

## Materials and Methods

### Subject selection

Subjects were selected by database searches of the Arizona Study of Aging and Neurodegenerative Disorders (AZSAND)/ Banner Sun Health Research Institute Brain and Body Donation Program (www.brainandbodydonationprogram.org) [93], a subset of whom were also enrolled in the National Institute on Aging Arizona Alzheimer’s Disease Core Center. Search criteria specified that subjects died with dementia, one or more completed University of Pennsylvania Smell Identification Tests (UPSIT) accompanied by Mini Mental State Examinations (MMSE), assessments of the presence or absence of parkinsonism and visual hallucinations, and a full neuropathological examination after death. Selected subjects met “intermediate” or “high” National Institute on Aging-Reagan Institute (NIA-RI) clinicopathological criteria [94] for ADD, with or without also meeting “intermediate” or “high” clinicopathological criteria for DLB [43,44], or alternatively, for a group termed as AD-LB [91], also having pathologically-confirmed CNS LBD but not meeting DLB pathology distribution and density thresholds. A control subject group, without clinical parkinsonism or dementia, and without α-synuclein pathology at autopsy, was also included. For all subjects, most other major neuropathological disorders were excluded; this included subjects with both ADD and Parkinson’s disease, progressive supranuclear palsy, multiple system atrophy and corticobasal degeneration. As mean UPSIT did not differ between ADD and the neuropathologically-defined AD-LB (n=30) and AD-VaD (n=25) groups, these were included with the ADD group for the primary analysis.

### Subject characterization

Subjects all had serial standardized research cognitive evaluations, done by teams of nurses, medical assistants, behavioral neurologists, movement disorders neurologists, neuropsychologists and psychometrists using standardized research-quality assessment batteries [93], including the Mini Mental State Examination (MMSE), National Alzheimer’s Coordinating Center (NACC) Uniform Data Set (UDS) and the Unified Parkinson’s Disease Rating Scale (UPDRS). Subjects had olfactory testing with the University of Pennsylvania Smell Identification Test (UPSIT)[95–97] every third year on average. The presence or absence of DLB core clinical features [43,44], including the presence or absence of parkinsonism, visual hallucinations, fluctuations in attention or cognition and clinical history consistent with REM sleep behavior disorder (RBD), were recorded for each subject at each visit; to assist with the latter, the Mayo Sleep Questionnaire [98–101] was administered. The presence of parkinsonism and visual hallucinations was additionally noted by review of private medical records and were used for comparison purposes as alternate predictors of DLB. These were determined within the same year as the first UPSIT administration, for comparison with first UPSIT as a diagnostic predictor, while for comparison with mean UPSIT score, the cumulative recorded presence of visual hallucinations and parkinsonism at any timepoint within the clinical observation period was used.

All subjects received identical neuropathological examinations, including summary regional brain density measures for total amyloid plaques, neurofibrillary tangles, Lewy body pathology regional and summary density scoring, and staging using the Unified Staging System for Lewy Body Disorders [91], as well as assignment of CERAD neuritic plaque density and Braak neurofibrillary stage, as described previously [93].

### Statistical analysis

Demographic and post-mortem characteristics were analyzed using one-way analysis of variance (ANOVA), Chi-square tests and unpaired t-tests as appropriate. Receiver-operator characteristics analysis was implemented for first and mean UPSIT scores to predict the diagnosis of DLB vs. ADD. Youden index was used as the criteria to choose the optimum cut-off point for UPSIT scores. Sensitivity, specificity, and accuracy of predicting DLB based on different UPSIT cutoff scores and the presence or absence of visual hallucinations and parkinsonism were further calculated. Firth logistic regression models adjusted for age, gender and corresponding MMSE scores were used to estimate odds ratios for different predictors and area under the curve (AUC) for each model. The AUCs for models with UPSIT, visual hallucinations and parkinsonism were compared using Delong’s method [102].

## Results

Clinical, demographic and neuropathological characteristics of the compared groups (total n = 209) are shown in Table 1. The DLB group (n = 29) had a significantly higher proportion of men and was slightly younger than the other two groups but this did not reach the significance level. As expected, the control group (n = 84) was significantly different from the two dementia groups in all clinical and neuropathological measures. The ADD (n = 96) and DLB groups were not significantly different on their last MMSE scores but the DLB group had significantly higher final motor UPDRS scores. Neuropathologically, the ADD and DLB groups were not different in their density scores or stages for amyloid plaques (total plaques), neuritic plaques and neurofibrillary tangles. As expected by group definition, Lewy body pathology brain load was much higher for the DLB group as compared to the ADD group.

**Table 1.**
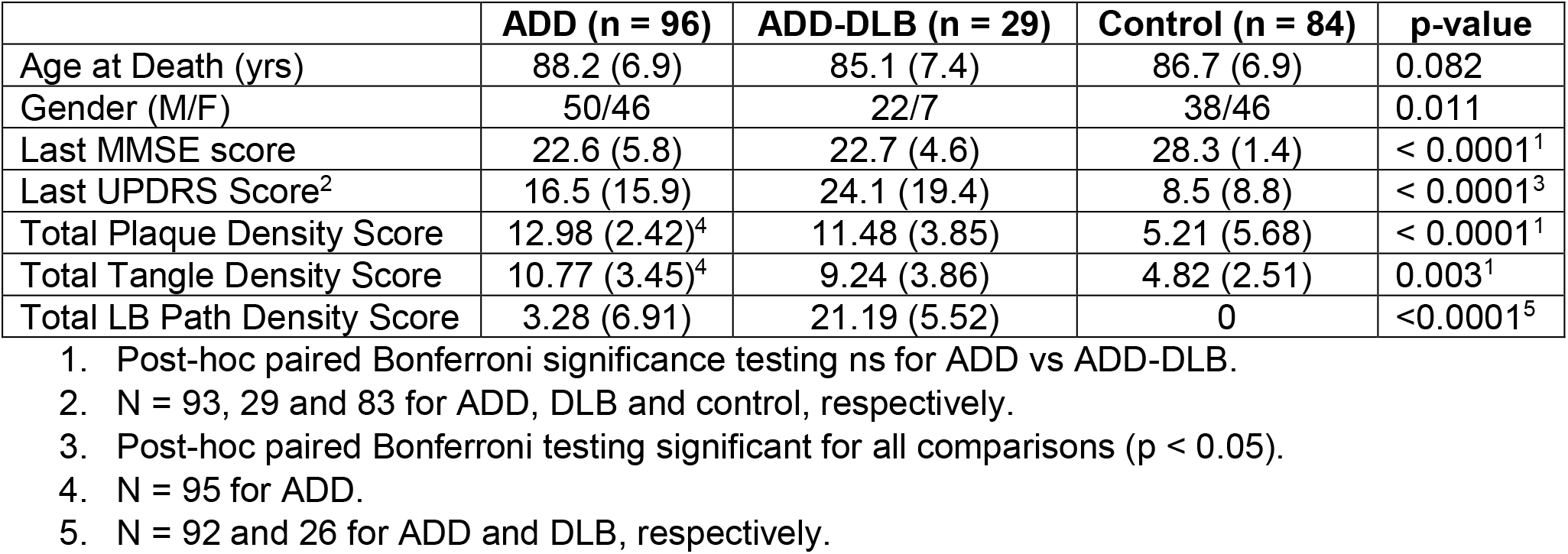
Clinical and neuropathological characteristics of study subjects. Means and standard deviations are shown. ADD = Alzheimer’s disease dementia; DLB = dementia with Lewy bodies; MMSE = Mini Mental State Examination score; UPDRS = Unified Parkinson’s Disease Rating Scale motor score (part 3 score); Total LB Path = total Lewy-type synucleinopathy density score

Comparison of UPSIT scores, including the first UPSIT score and the mean of all UPSIT scores, showed that the DLB group had significantly lower scores than both other groups (Figure 1). Control subjects had significantly more UPSITs (mean 1.9, range 1-4; p < 0.05) than ADD (mean 1.6, range 1-4) or DLB subjects (mean 1.35, range 1-3) but this did not differ between ADD and DLB groups. The time intervals between first UPSIT and death did not differ between the two dementia groups; for ADD and DLB these were 6.1 years and 5.0 years, respectively. Logistic regression analysis of the combined DLB, ADD and control groups (n = 188 after exclusion of cases with incomplete neuropathology scores) found only the brain regional sum of α-synuclein pathology scores (maximum score of 40) was significantly associated with an UPSIT score less than the median of all 188 cases (OR 1.14, 95% CI 1.06-1.22, p = 0.0002)). Total regional brain scores (maximum of 15) for amyloid plaque and neurofibrillary tangle density, as well as age at death, were not significant predictors, while last MMSE test score approached the significance level (OR 0.96, 95% CI 0.91-1.01, p = 0.098).

**Figure 1.**
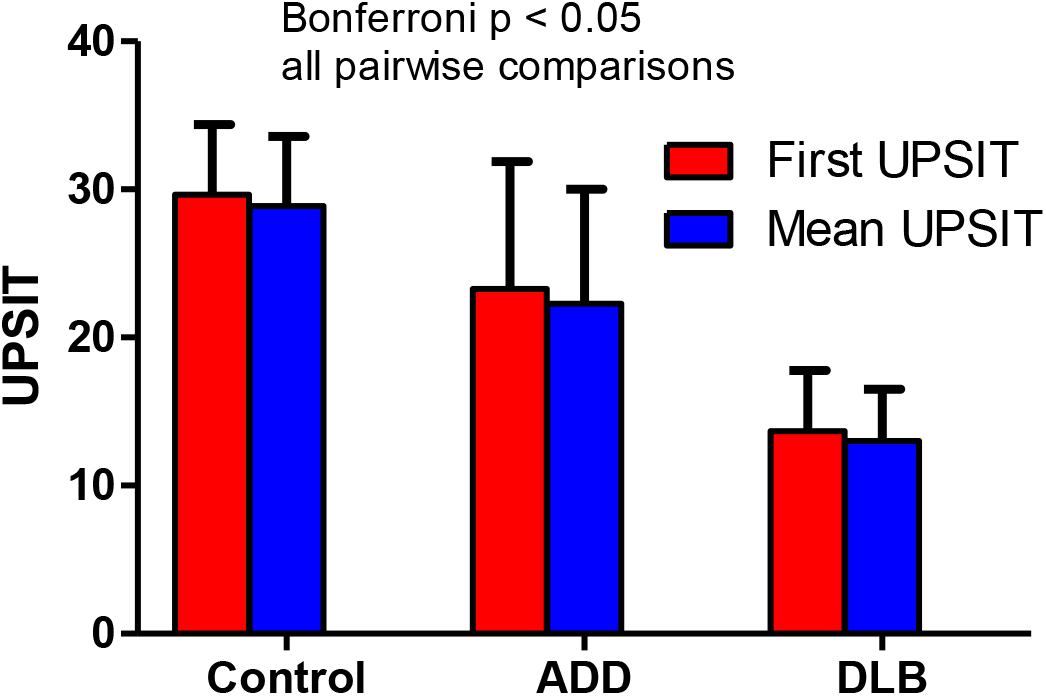
First and mean UPSIT scores in the three diagnostic groups. Both first UPSIT and mean UPSIT scores are significantly different between groups (ANOVA, p < 0.001) and all pairwise group comparisons are significant (Bonferroni p < 0.05). First and mean UPSIT scores were not significantly different within diagnostic groups. Error bars = standard deviation.

Logistic regression analysis and receiver-operator characteristics (ROC) curves indicated that the UPSIT cutoff scores giving the greatest accuracy for separating ADD from DLB were 19 or less for the first UPSIT (median of DLB and ADD first UPSIT scores was 20) and 17 for the mean UPSIT. Using these cutoffs, a first UPSIT score less than 20 gave an odds ratio (OR) of 17.5 for a diagnosis of DLB, while for mean UPSIT, a score less than 17 resulted in an OR of 18.0 for the diagnosis (Table 2). These ORs were considerably greater than those derived from the presence or absence of the two most common DLB core clinical features, visual hallucinations and parkinsonism (Table 2, Figure 2a,b), and were highly significant (p < 0.0001) whereas only the OR for cumulatively-observed hallucinations was significant (OR 3.3; p = 0.01). Similarly, the area under the curves (AUC) were significantly greater for first and mean UPSIT as compared with those for matched-year or cumulative hallucinations and parkinsonism.

**Table 2.** Comparison of first UPSIT score and mean of all UPSIT scores with visual hallucinations and parkinsonism as predictors of DLB.

| Predictor | Sensitivity | Specificity | Accuracy | Odds Ratio (95% CI), p-value | AUC | P-value |
| --- | --- | --- | --- | --- | --- | --- |
| First UPSIT<sup>1</sup> | 93.1% | 64.6% | 71.2% | 17.5 (5.1, 91.6)<br><.0001 | 82.9% | 0.2419<sup>3</sup> |
| Matched Year hallucinations<sup>1</sup> | 17.2% | 96.9% | 78.4% | 4.4 (0.9, 25.0)<br>0.0905 | 67.4% | 0.0012<sup>4</sup> |
| Matched Year parkinsonism<sup>1</sup> | 31.0% | 77.1% | 66.4% | 1.7 (0.7, 4.3)<br>0.2648 | 67.4% | 0.0006<sup>4</sup> |
| Mean UPSIT<sup>2</sup> | 86.2% | 71.9% | 75.2% | 18.0 (6.0, 66.8)<br><.0001 | 87.2% | 0.2419<sup>3</sup> |
| Cumulative hallucinations<sup>2</sup> | 51.7% | 76.0% | 70.4% | 3.3 (1.4, 8.4)<br>0.0106 | 72.9% | 0.008<sup>4</sup> |
| Cumulative parkinsonism<sup>2</sup> | 65.5% | 45.8% | 50.4% | 1.6 (0.7, 3.9)<br>0.3001 | 68.2% | 0.0007<sup>4</sup> |

1. Adjusted for matched year MMSE and age at first UPSIT 2. Adjusted for mean MMSE and age at death 3. P-value comparing AUCs for first and mean UPSIT 4. P-value comparing AUC with first UPSIT 5. P-value comparing AUC with mean UPSIT

**Figure 2.**
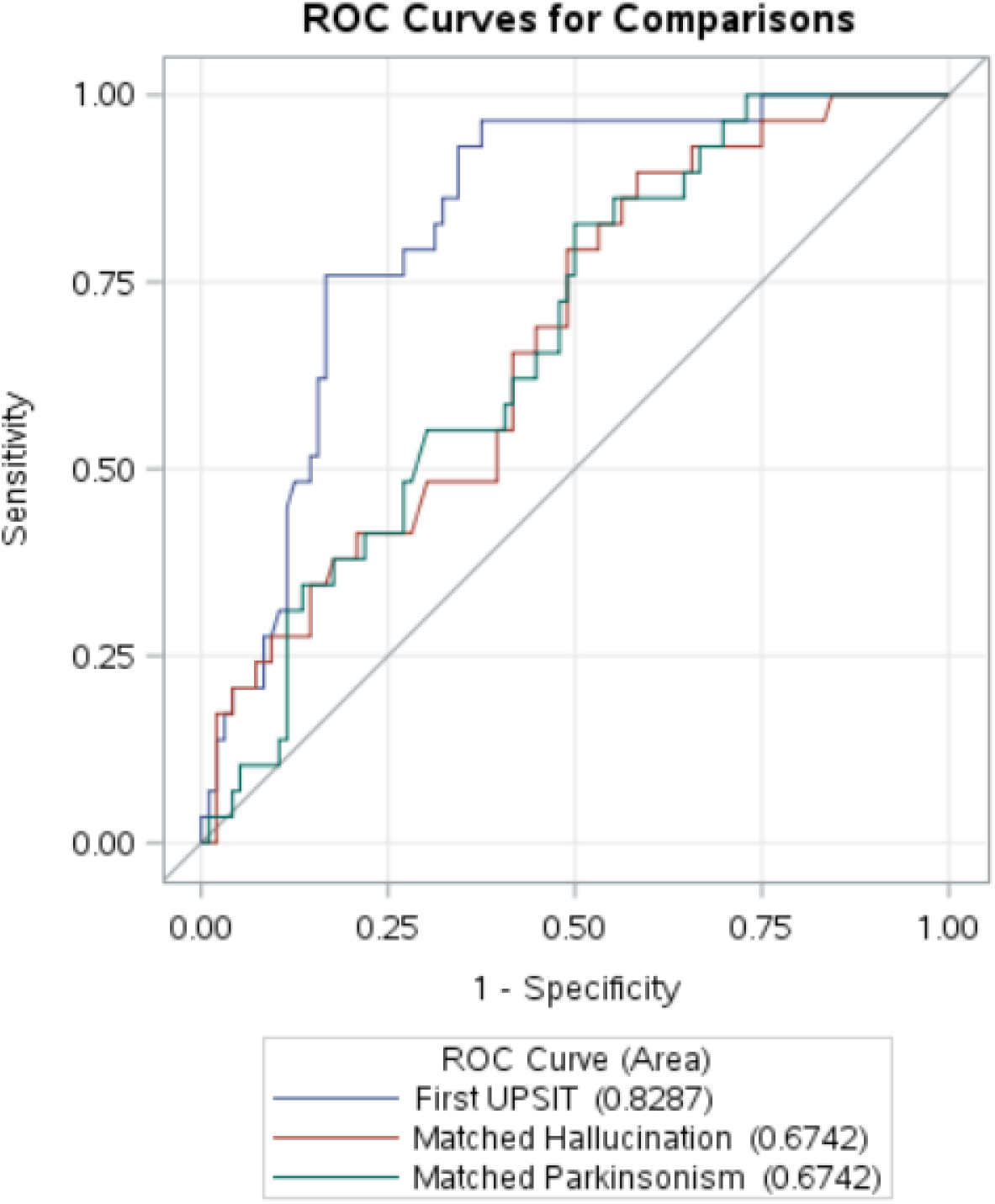
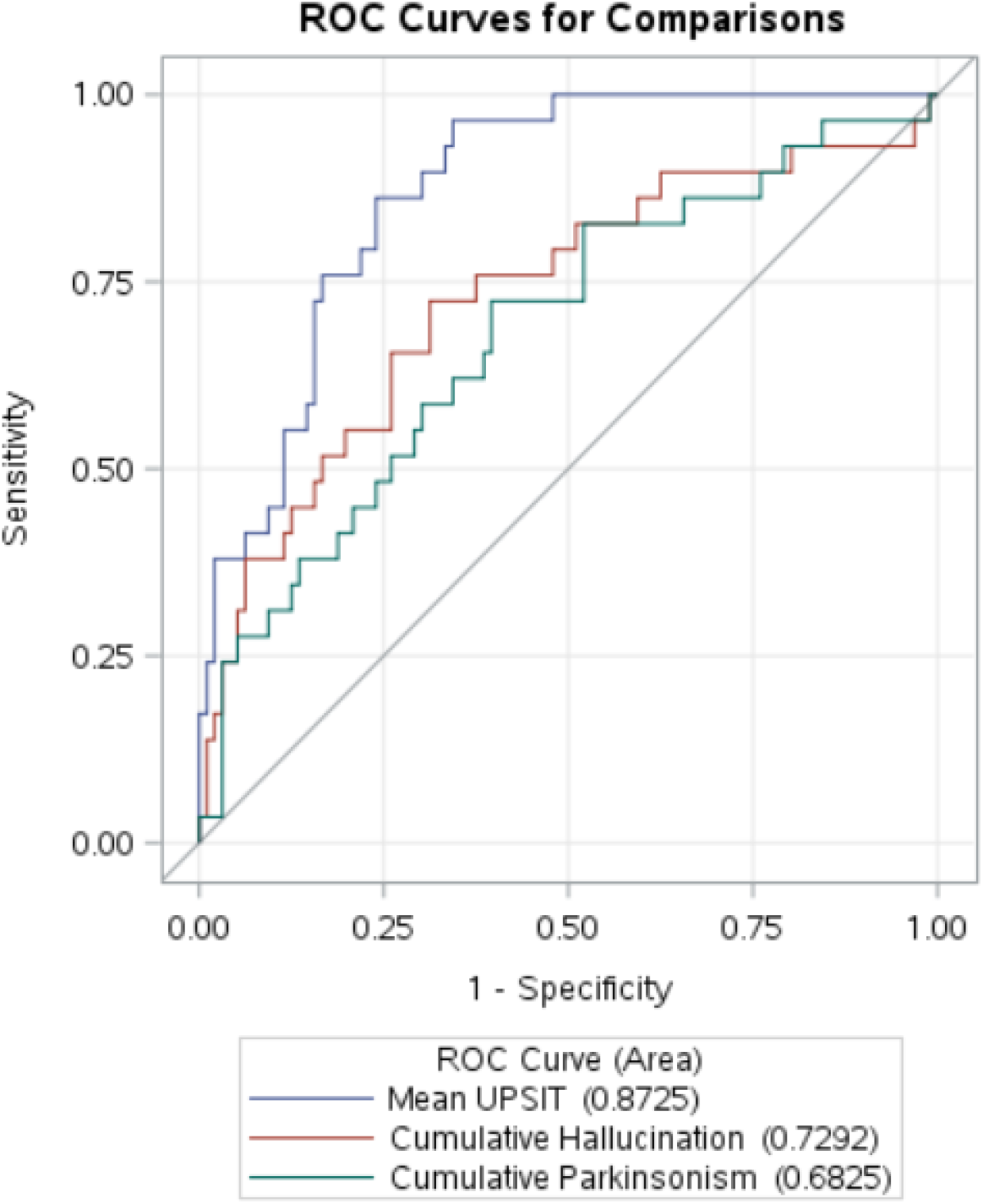
**a-b.** Comparison of ROC curves for a) first UPSIT scores and b) mean UPSIT scores, with those for presence or absence of visual hallucinations and parkinsonism.

To determine whether the inclusion of ADLB subjects with the ADD group affected the primary analysis, logistic regression analysis and ROC curves were used to compare the ADD and DLB groups after exclusion of the 30 subjects with ADLB. The results were very similar to those obtained with the ADD and ADLB groups together (Figure 3a, 3b). All comparisons that were statistically significant in the primary analysis were also significant after exclusion of the ADLB subjects. A first UPSIT cutoff score of 19 gave an OR of 28.3 for separating ADD from DLB while a mean UPSIT cutoff of 20 gave an OR of 24.4 for predicting DLB.

**Figure 3.**
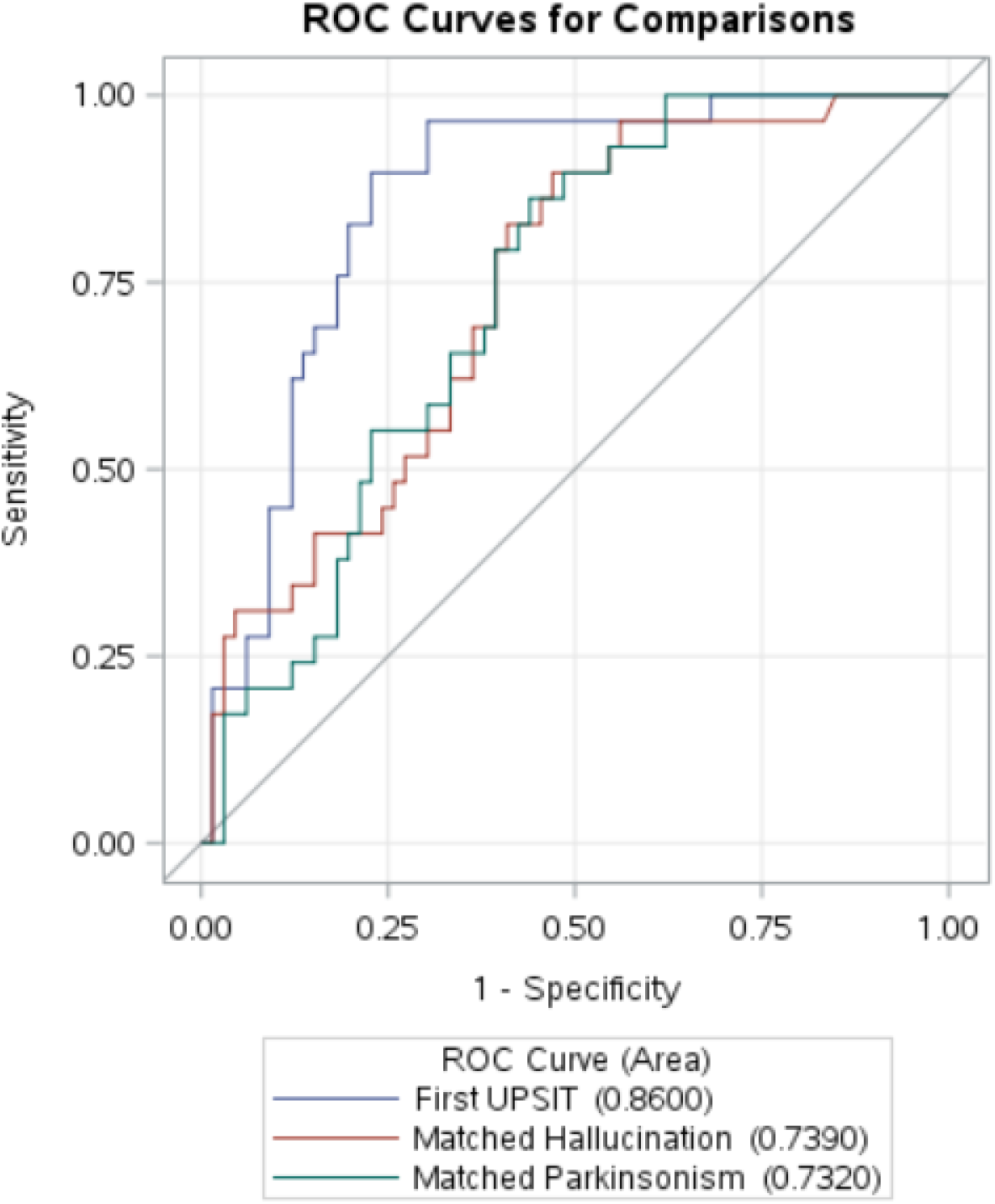
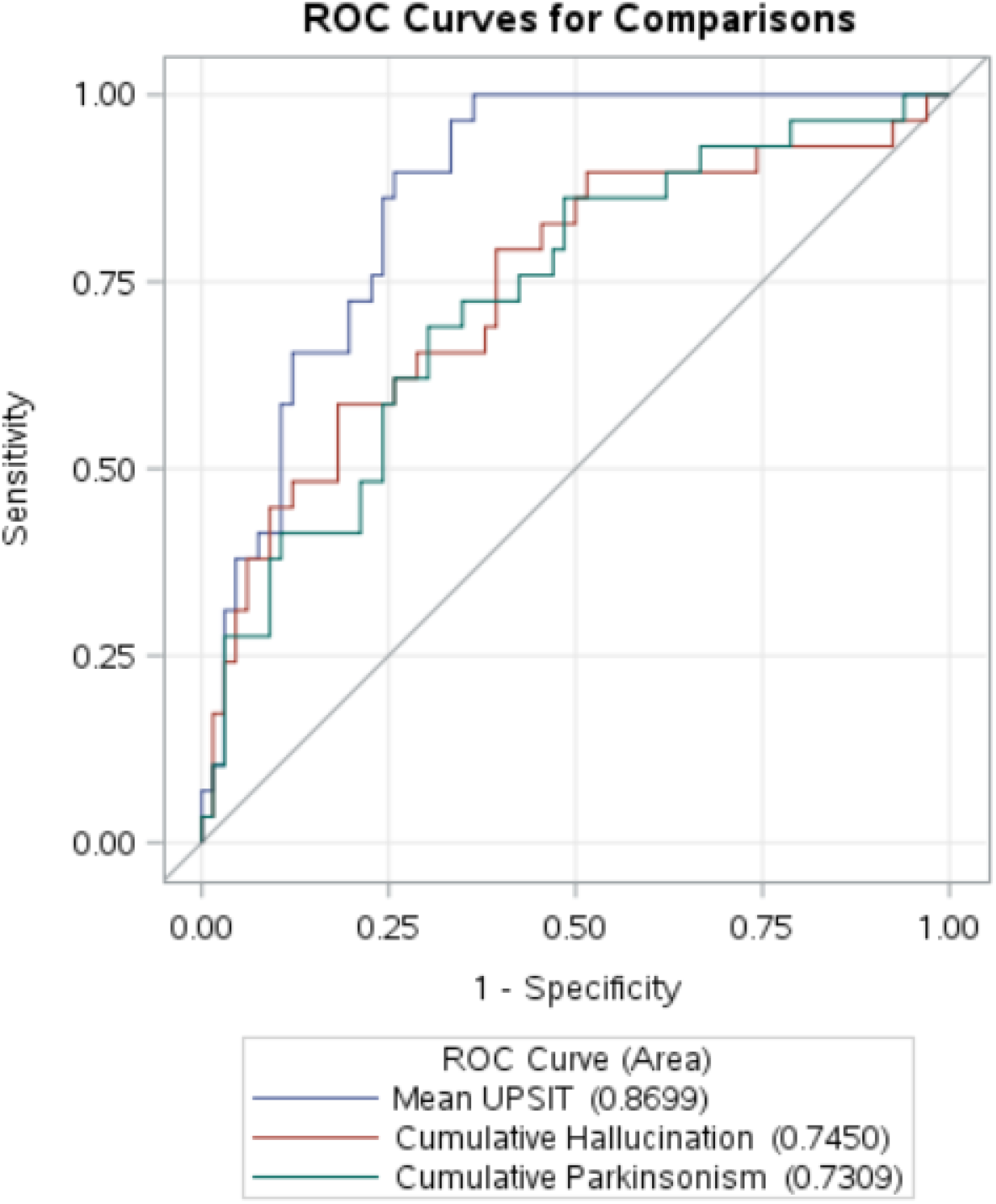
**a-b.** Comparison of ROC curves after exclusion of 30 ADLB subjects from the ADD group. a) first UPSIT scores and b) mean UPSIT scores.

## Discussion

ADD and DLB often co-exist unknown to clinicians, and as this comorbidity may affect the presentation, rate of cognitive decline, and response to therapeutic agents [1–22], clinical trials for both conditions may be impaired. The most common comorbidity in ADD is LBD, affecting somewhat more than one-half or more of all those meeting clinicopathological ADD diagnostic criteria [9,23–25]. Dementia with Lewy bodies has greater α-synuclein pathology than ADLB and therefore may be more resistant to therapeutic agents targeting AD molecular lesions. In the majority of subjects with ADD and DLB, the typical clinical signs and symptoms of DLB [43,44] are absent and thus this co-existence is recognized only at autopsy [22,45–47]. Disease duration has been reported to be shorter in those with coexistent ADD and DLB [40,52]. There is therefore a critical need for better clinical differentiation of these two conditions.

Definitive laboratory-based biomarkers for DLB are not yet available. Functional dopamine imaging and myocardial scintigraphy with [123I]meta-iodobenzylguanidine (MIBG) have both been used as diagnostic adjuncts for DLB [103,104] with promising but not yet definitive results from small autopsy-confirmed studies [105,106]. Dopaminergic imaging may be less helpful in DLB as compared to PD, due to lesser loss of nigrostriatal dopaminergic neuronal and nerve terminals [107–111]. Biofluids and PET imaging approaches have so far been unsuccessful in providing the required accuracy for identifying LBD [112–114]. Simulation studies have suggested that cortical biopsy [115–118] would have high sensitivity and specificity for DLB, and usage of needle cores rather than open biopsy may reduce morbidity to acceptable levels [115]. Biopsy of the peripheral nervous system [119], particularly the submandibular gland [120–122], also shows promise for diagnosing DLB. Autopsy studies have suggested that biopsy of the olfactory bulb would identify more than 90% of all subjects with LBD [90]. Better clinical diagnostic methods for DLB are critically needed, as improved sensitivity in the clinical identification of DLB would greatly assist recruitment for clinical trials and would allow exclusion or stratification of DLB subjects within ADD clinical trials.

Numerous studies have indicated the potential for olfactory function tests to distinguish different cerebrovascular and neurodegenerative disorders [53–64] and, in particular, to distinguish PD and DLB from ADD [65–70], but the great majority of these studies lack certainty due the reliance on a clinical diagnosis as gold standard. Several studies with neuropathological confirmation of LBD have suggested that loss of olfactory function may be more pronounced in DLB but these have been limited by small subject numbers [71–74].

In this study we sought to determine the diagnostic utility of hyposmia as a diagnostic predictor of neuropathologically-identified DLB, using considerably larger subject numbers than previous studies. Our results confirm those of the prior studies, where subjects with DLB have been repeatedly found to have worse olfactory function than ADD. The odds ratios for ROC-determined first and mean UPSIT score cutoffs, 17.5 and 18.0, respectively, were surprisingly stronger than the ORs for both visual hallucinations and parkinsonism (1.7 - 4.4), two of the key core clinical DLB features. These figures suggest that olfactory testing should be considered as a core clinical feature of DLB and could potentially be of great assistance in the clinical separation of ADD and DLB, allowing stratification of clinical trial subjects. Larger neuropathologically-examined subject numbers would help to confirm the results of the present study but if results from the prior three neuropathologically-confirmed studies are added to this, there are 89 LBD cases and 365 controls (ADD or normal controls), all with the same general finding of much lower olfactory test scores in the LBD groups.

As the first and mean UPSIT scores were not significantly different, it seems probable that hyposmia is an early clinical occurrence in DLB, and hence smell testing could be helpful in the identification of prodromal DLB. Support for this possibility comes from studies of incidental Lewy body disease (ILBD), defined as the presence of LBD in asymptomatic elderly people. ILBD is a probable prodromal stage of PD or DLB as dopaminergic markers are halfway between asymptomatic elderly people without LBD and clinically-manifest PD [75–78]. Our group has previously reported that olfactory function in subjects with ILBD is also halfway between PD and asymptomatic elderly people without LBD [79]; another clinicopathological study found an OR of 11.0 for hyposmia in the prediction of ILBD, using as a cutoff the lowest tertile of olfactory function [80], and these postmortem studies have been further confirmed by the in vivo association of hyposmia with decreased striatal dopamine transporter imaging [87,88]. Additional support comes from reports of hyposmia in some clinically normal GBA and LRRK2 mutation carriers [81,82] and in idiopathic REM sleep behavior disorder (iRBD) [83–86].

## Acknowledgements and Funding

The Banner Sun Health Research Institute Brain and Body Donation Program has been supported by the National Institute of Neurological Disorders and Stroke (U24 NS072026 National Brain and Tissue Resource for Parkinson’s Disease and Related Disorders), the National Institute on Aging (P30 AG19610 Arizona Alzheimer’s Disease Core Center), the Arizona Department of Health Services (contract 211002, Arizona Alzheimer’s Research Center), the Arizona Biomedical Research Commission (contracts 4001, 0011, 05-901 and 1001 to the Arizona Parkinson’s Disease Consortium), Sun Health Foundation, Mayo Clinic Foundation, and the Michael J. Fox Foundation for Parkinson’s Research.

